# Desensitization, Inactivation, and the tension-proof safety mechanism of inactivated MscS

**DOI:** 10.64898/2026.03.12.711414

**Authors:** Andriy Anishkin, Elissa Moller, Sergei Sukharev

## Abstract

MscS is the main low-threshold tension-activated osmolyte release valve in bacteria. Working alone or together with high-threshold MscL, it regulates turgor and protects cells from mechanical rupture during osmotic down-shock. The channel exhibits complex adaptive behavior, including desensitization and full inactivation, both of which occur at relatively low sub-lytic tension. There is debate over whether the commonly observed non-conductive state of MscS with the tension-sensing helices splayed away from the gate corresponds to the closed or inactivated state. In this work, using specialized pressure protocols in patch-clamp electrophysiology, we highlight the difference between reversible adaptation (desensitization) and inactivation. We show that inactivated channels cannot be reactivated with high tension, up to the limit of patch stability. This aligns with cryo-EM studies by Zhang et al. (*Nature*, 2021, 590:509-5018), who applied extreme tension to the splayed nanodisc-reconstituted MscS (PDB 6VYK) by depleting lipids with cyclodextrin, and observed a new flattened but apparently non-conductive structure (PDB 6VYM). To characterize these two states, we performed a steered Molecular Dynamics simulation from the initial splayed structure to the flattened conformation, confirming that they are connected through a smooth conformational pathway and remain largely dehydrated and entirely non-conductive throughout the transition. The data show that the initial splayed conformation meets all the criteria of the inactivated state, distorting but not opening under extreme tension. By combining patch-clamp experiments with simulations based on cryo-EM data, we demonstrate that inactivated MscS resists activation, thereby maintaining the membrane barrier when tension exceeds the activation threshold.

**SIGNIFICANCE:** Bacterial energetics, which relies on the electrochemical proton gradient across the inner membrane as an intermediary, conflicts with the presence of a dense population of highly conductive mechano-activated channels in the same cytoplasmic membrane, which must be proton-tight. Moreover, the tension activation threshold for the common MscS channel is low and can be easily exceeded by fluctuations in the concentrations of internal or external osmolytes. In this paper, we describe the important adaptive inactivation of MscS at tensions near its activation threshold, and specifically, the complete resistance of inactivated MscS to opening at any tension. The data show another layer of tight regulation of MscS residing in the energy-coupling membranes.

## INTRODUCTION

As the main component of the turgor-adjustment system in prokaryotes (1,2), MscS enables rapid bacterial adaptation to freshwater, the main transmission route for most commensals and pathogens (3–6). It resides in the cytoplasmic membrane and acts as an adaptive osmolyte release valve activated by sub-lytic tensions typically generated by moderate osmotic downshifts. In *Escherichia coli*, MscS works in conjunction with the high-threshold channel MscL, which activates at near-lytic tensions, thus forming a release system that provides a graded permeability response to low, moderate, and extreme osmotic shocks (7,8). Some species lack MscL (9), but one or more genes encoding MscS-like channels are always present in microorganisms that spend at least part of their life cycle in the environment. MscS perfectly rescues bacterial cells from osmotic damage in the absence of MscL (2,10).

The osmo-protective mechanism of bacterial mechanosensitive channels is kinetic (2,10,11). When under strongly hypotonic conditions, water rapidly swells the cell, the counteracting release of osmolytes must outpace the influx of water. Kinetic measurements showed that under severe down-shock, cells can release up to 15% of their dry weight as small metabolites within 10-20 ms (2,11). Estimates show that during the release phase, membrane porosity must increase by at least 10 orders of magnitude by opening hundreds of membrane-embedded channels per cell. The question arises: how safe is it to have large populations of osmolyte release valves in the energy-coupling bacterial membrane? Indeed, the chemiosmotic mechanism coupling oxidative respiration and ATP synthesis is mediated by the electrochemical proton gradient (12), which can be shunted by highly conductive, essentially non-selective channels in the energy-coupling membrane. A spurious opening of a single MscS channel (1 nS unitary conductance) will deenergize the bacterial cell within 0.2 ms. The only way to reconcile the presence of channels with the membrane’s energy-coupling function was to assume their tight regulation (13).

When the MscL and MscS channels were cloned (1,14) and structurally characterized (15,16), it became clear that, despite large conformational changes, their hydrophobic gates are dehydrated (vapor-locked) at rest and should be leak-proof (17,18). While MscL, with its midpoint of 12-14 mN/m, must be firmly closed under most physiological conditions (19,20), the low-threshold MscS showed activity at moderate 5-7 mN/m (21,22). The question was whether these tensions could fall within the normal range in an actively growing cell at any point in its life cycle in an osmotically balanced medium. A partial answer came with the discovery of MscS inactivation, a transition to a non-conductive, tension-insensitive state driven by low, near-threshold tension from the closed state (23–25). This slow transition is expected to prevent spurious channel activity when tension fluctuates near a non-threatening level of 5-6 mN/m for a prolonged period. In our recent structure (PDB 9P0N), we demonstrated that lipid penetration into interhelical crevices uncouples the tension-receiving TM1-TM2 pairs from the gate, rendering the channel tension-insensitive (26). The next question is how reliable this uncoupling mechanism is and whether stronger tensions could potentially reactivate the channel.

The recent study by Zhang and coworkers (27) provided critical structural insights into the transformation of the non-conductive MscS conformation with lipid-filled crevices (PDB 6VYK), labeled ‘closed’, into a highly expanded state (6VYM), observed under extreme tension and labeled ‘desensitized’. The cryo-EM structures were determined in PC18:1 nanodiscs; the initial splayed state was solved in unperturbed reconstituted particles, whereas the expanded state was obtained after a prolonged exposure of the nanodiscs to Methyl-β-cyclodextrin. According to their estimates, cyclodextrin depletes lipids from nanodiscs by about 35%, generating an internal tension of 30-40 mN/m (extended data Fig. 7 (27)), well beyond the stability limit of all known bilayers. Despite the highly unphysiological conditions, the properties of this expanded and flattened structure, as well as the transition leading to it, warrant further characterization.

In this work, we reinterpret the structures published by Zhang and colleagues and classify the splayed structure with peripheral helices decoupled from the gate and a 6.1 Å hydrophobic pore (PDB 6VYK) as inactivated (27). We simulate the transition from the inactivated state to the expanded (PDB 6VYM) conformation in an E. coli-like bilayer and find that its 6.5 Å pore remains partially de-solvated and completely non-conductive. In patch-clamp experiments, we emphasize the distinction between desensitized and inactivated states and demonstrate that the inactivated MscS cannot be reopened at any tension attainable in a typical patch-clamp setting. We present the conformational connectivity between the two experimental structures and interpret the flattened 6VYM structure as a highly distorted inactivated state. The data illustrate that the functional design of MscS is exceptionally robust and ‘safe’. Once inactivated, the channel remains non-conductive not only above the activation threshold but also at extreme tensions outside the physiological range.

## MATERIALS AND METHODS

### Patch-clamp experiments

The MscS channel was expressed from the pB10d plasmid (28) in the MJF465 (ΔMscL ΔMscS ΔMscK) *E. coli* strain (1). Giant spheroplasts were generated as described previously (29) by growing filamentous forms in the presence of 0.06 mg/ml cephalexin for 1.5 hrs. Protein expression was induced with 1 mM IPTG for the final 30 min. The resulting filaments were transferred into a 1 M sucrose with 1.8 mg/ml BSA buffer and treated with 0.17 mg/ml lysozyme in the presence of 4 mM EDTA (8 min), resulting in the formation of round cells up to 4-5 mm in diameter (giant spheroplasts). The giant spheroplasts were isolated by sedimentation through a one-step sucrose gradient (1.2 M).

Patch-clamp traces were recorded from populations of 100-150 channels in excised inside-out patches at room temperature. The recording buffers contained 200 mM KCl, 50 mM MgCl_2_, 5 mM CaCl_2_, and 5 mM HEPES at pH 7.4 on both sides, whereas the bath solution contained an extra 400 mM sucrose to osmotically stabilize the spheroplasts. Membrane patches were obtained using borosilicate glass pipettes (Drummond) and recorded on an Axopatch 200B amplifier (Molecular Devices). Negative pressure stimuli were applied using a high-speed pressure clamp apparatus (HSPC-1; ALA Scientific) programmed using the Clampex software (Molecular Devices). Pressure and voltage protocol programming and data acquisition were performed with the PClamp 10 suite (Axon Instruments). The pressure protocols designed to discern between adapted and inactivated states are described in the text.

### Molecular Dynamics Simulations

The simulation system was assembled using the CHARMM GUI 71 around the 6PWP cryo-EM structure (Reddy et al., 2019), which resolved the most complete sequence, and was then converted to 6VYK (27). The lipid bilayer, composed of 498 POPE and 166 POPG molecules (ratio 3/1), was assembled around the channel to bring the lateral size of the orthogonal simulation cell to 150 x 150 Å, allowing at least 9-10 lipid layers between the closest mirror images of the protein. The membrane protein system was hydrated, and K+ and Cl-ions were added to make a 200 mM KCl salt solution, bringing the total number of atoms in the system to ~395,000. The entire system was energy-minimized with a restrained protein backbone for 10,000 steps using the conjugate energy gradient algorithm, and then simulated for 20 ns with the protein backbone harmonically restrained near the modeled positions using a spring constant of 1 kcal/mol/Å.

Because the 6VYK structure was resolved beginning with Tryptophan 16 (27), the restraints of the simulated 6PWP were released for the N-terminal segment, from residue 1 through 16, while the backbone of all resolved residues was pulled toward 6VYK, thus converting one structure into another. Although the backbone RMSD between the transmembrane domains (residues 15 to 128) of 6PWP and 6VYK is only 2.1 Å, and the adjustment was minor, the obtained 6VYK with liberated N-termini was additionally equilibrated for 20 ns with the backbone restrained at 1 kcal/mol/A^2^.

All simulations were performed in a flexible, orthogonal cell under periodic boundary conditions, at 1 bar pressure, and with a lateral tension of 20 dyne/cm, using NAMD2 (30,31) with the CHARMM27 force field (32) and the TIP3P water model (33). Particle mesh Ewald method was used for long-range electrostatics estimation 74, a 10 Å cutoff for short-range electrostatic and Van der Waals forces, and a Langevin thermostat set at 310 K. After the restrained backbone stage was completed, the system was simulated unrestrained for 100 ns, with the last 80 ns used to quantify the contact statistics. Visualization of the channel structures was performed using Visual Molecular Dynamics (VMD) (34). All structural and statistical analyses for MD simulations were performed using custom Tcl scripts in VMD.

To explore the conformational connectivity between the structures 6VYK and the flattened 6VYM (27), we steered this 100 ns transition using the Targeted Molecular Dynamics module in NAMD. To allow side chains freedom to move and adjust to the membrane, we steered only the backbone of residues 28-279 (all that were clearly resolved in the 6VYM structure). The force was applied uniformly to all steered atoms, pushing them toward their respective target coordinates. The magnitude of the uniform force was based on harmonic restraints with a spring constant of 1 kcal/mol/Å^2^. During steering, the force was adjusted up or down to approximate a linear decrease in RMSD from the starting value to the target (zero). For this, at each timestep, the constant was multiplied by the difference between the current RMSD of all steered atoms and the expected RMSD for a linear decrease. Because the same force was applied to each steered atom, we plotted the force per atom to illustrate the character of the transition and infer possible energy barriers along the way.

The water pore occupancy was calculated within a 5 Å slab across the pore (in the plane of the membrane), centered on the alpha-carbons of L105 and L109, which form the hydrophobic gate, and plotted against simulation time. The time-averaged distribution of water density along the pore was estimated from a coaxial slice of the volumetric density map of water oxygens. For each atom, its mass was distributed uniformly within a sphere with the CHARMM36 VDW radius for water oxygen. The volumetric distribution was averaged over the entire 100 ns of the simulation with a restrained backbone. The map was color-coded from red (vacuum) to blue (bulk density of water oxygens). Because we were interested in detecting permeant ions in the initial and end states as well as along the transition, all simulations were performed under a constant cytoplasmic voltage of −200 mV. Volumetric maps of Cl^−^ ion distribution were calculated using VMD.

## RESULTS

### Sequential adaptive closure and inactivation of MscS

Inactivated channels cannot be reactivated at any tension without a recovery period

The four traces in Fig. 1 (A-D) demonstrate the key functional features of *E. coli* MscS, recorded on a single patch for consistency. The responses to symmetrical 1-second ramp stimuli (panel A) show the tension-activation curve for a population of approximately 100 channels. During the 1-second symmetric pressure ramp response, the midpoints of the rising and falling parts of the ramp are visibly different because of the differing tension dependence of the opening and closing rates (25,35). On the rising side, the first channel opened at −100 mm Hg, the midpoint was observed at −120 mm Hg, and the current saturated at −150 mm Hg in the pipette. Converting pipette pressure to membrane tension requires knowledge of patch curvature, typically obtained from patch or spheroplast imaging (19,21,36). Past measurements in whole-spheroplast mode showed that the midpoint of MscS activation is around 7.8 mN/m (36). Whole spheroplasts retain parts of the peptidoglycan and outer membrane, which can contribute to the envelope’s stiffness when the spheroplast is stretched by positive pipette pressure. This may lead to an overestimation of the midpoint in excised patches where the outer membrane and peptidoglycan are absent. In reconstituted liposome experiments using pure lipid patch imaging, MscS activates at a midpoint near 6 mN/m (21,22). It’s reasonable to assume that in excised spheroplast patches, which are protrusions of bare cytoplasmic membrane into the pipette, the midpoint is near 7 mN/m. Assuming the patch curvature stays constant, the saturating tension is 10 mN/m for this patch.

**Fig. 1.**
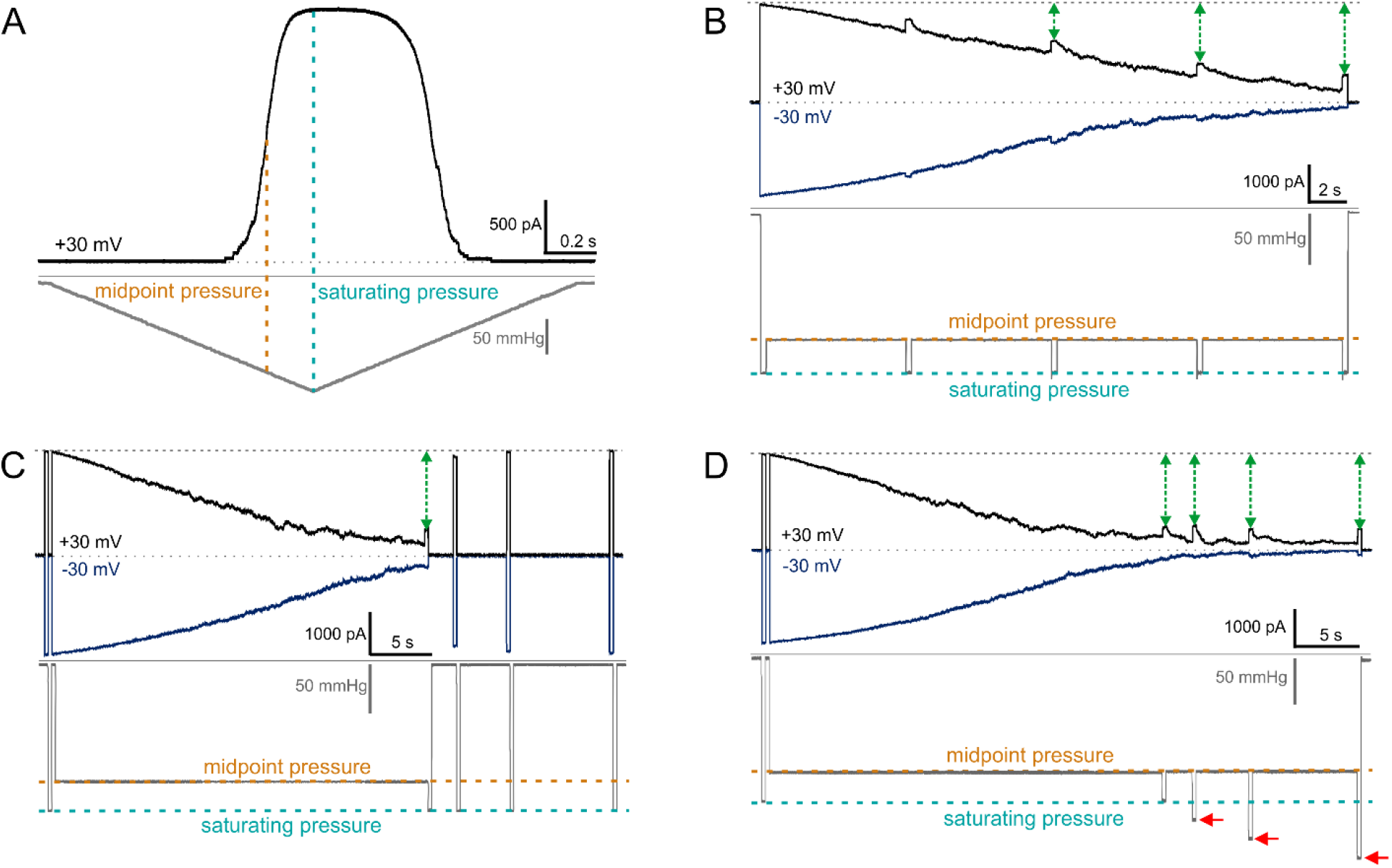
General phenomenology of MscS gating and adaptation revealed by four pressure protocols in an excised spheroplast patch containing about 100 MscS channels. (A) The ramp response reveals the midpoint and the saturation pressures. (B) The ‘comb’ protocol illustrates two parallel processes: gradual, reversible desensitization and silent inactivation from the desensitized closed state. (C) The inactivated population can fully reactivate after a 2-3-second rest at zero tension. (D) Increased amplitude of saturating test pulses (red arrows) at the end of the conditioning step does not activate any additional channels. Green arrows indicate the inactivated fraction of channels in panels B, C, and D. This phenomenology was observed in four patches from two separate batches of spheroplasts.

Panel B shows responses to the ‘comb’ pressure protocol, designed to distinguish between desensitization (adaptation) and complete inactivation. The protocol begins with a short, saturating pressure pulse (10 mN/m, 200 ms) that activates the entire population. The intermediate conditioning step then applies 7 mN/m tension for 30 seconds, matching the midpoint of the ramp response. Applying this pressure as an abrupt step, rather than a ramp, initially activates the entire population, which then begins to desensitize. Interspersed saturating pulses reactivate part of the reversibly adapted population, which gradually declines. The remaining part that fails to open (shown by the green two-headed arrows) belongs to the inactivated pool, which increases with more prolonged exposure to the midpoint tension.

Panel C shows how the same patch responds to a 30-second conditioning step, with two saturating test pulses on either side. The change in current during these test pulses indicates that by the 30th second, roughly 75% of the channels are inactivated. As seen from the rest of the trace, the inactivated channels can be reactivated after 2 seconds of rest at zero tension. Panel D demonstrates that when tension is held at the midpoint, increasing the saturating pulse amplitude by 50% (to 15 mN/m, the maximum limit of the pressure clamp machine) does not reactivate any additional channels. Under these conditions, we often observed patch rupture without reactivation of the inactivated channels.

This phenomenology of MscS gating and adaptive behavior observed with these pressure protocols shows a clear distinction between desensitized and inactivated states. The low-threshold channel MscS desensitizes first, then silently inactivates at tensions between 5 and 7 mN/m, i.e., between the threshold and the midpoint for activation. These tensions are not lethal or extreme and do not imply any thinning of the lipid bilayer surrounding the channel.

In the experiment presented in Fig. 2, we asked an additional question: how early in tension can we observe MscS inactivation? This series of traces was obtained in a separate patch, using ‘comb’ protocols similar to those in Fig. 1B. Evenly spaced saturating test pulses were applied on top of a prolonged conditioning step, which was varied. The midpoint pressure (corresponding to 7 mN/m tension, orange trace) determined in a ramp experiment produced a maximum of inactivation (95%) at the end of the 30 s protocol. Increasing or decreasing the amplitude of the conditioning step decreased the extent of inactivation. Increasing the step amplitude slowed adaptive closure. Because only closed channels can be inactivated (24), a smaller fraction of closed channels diminished the overall degree of inactivation. However, at high amplitudes of the conditioning step, all closed channels immediately inactivate, as indicated by almost no increase in current in response to saturating test pulses (green trace). Decreasing the amplitude of the conditioning step below the midpoint hastened adaptive closure (magenta, purple, and blue traces) but decreased the rate of inactivation. As a result, by the 30th second, the overall extent of inactivation was low. Yet at a tension of 5 N/m (blue trace), which corresponds to the activation threshold, we still observe ~10% inactivation. This illustrates that the slow inactivation process is driven by low tensions, and that the threshold for inactivation can be near or even slightly below that for activation.

**Fig. 2.**
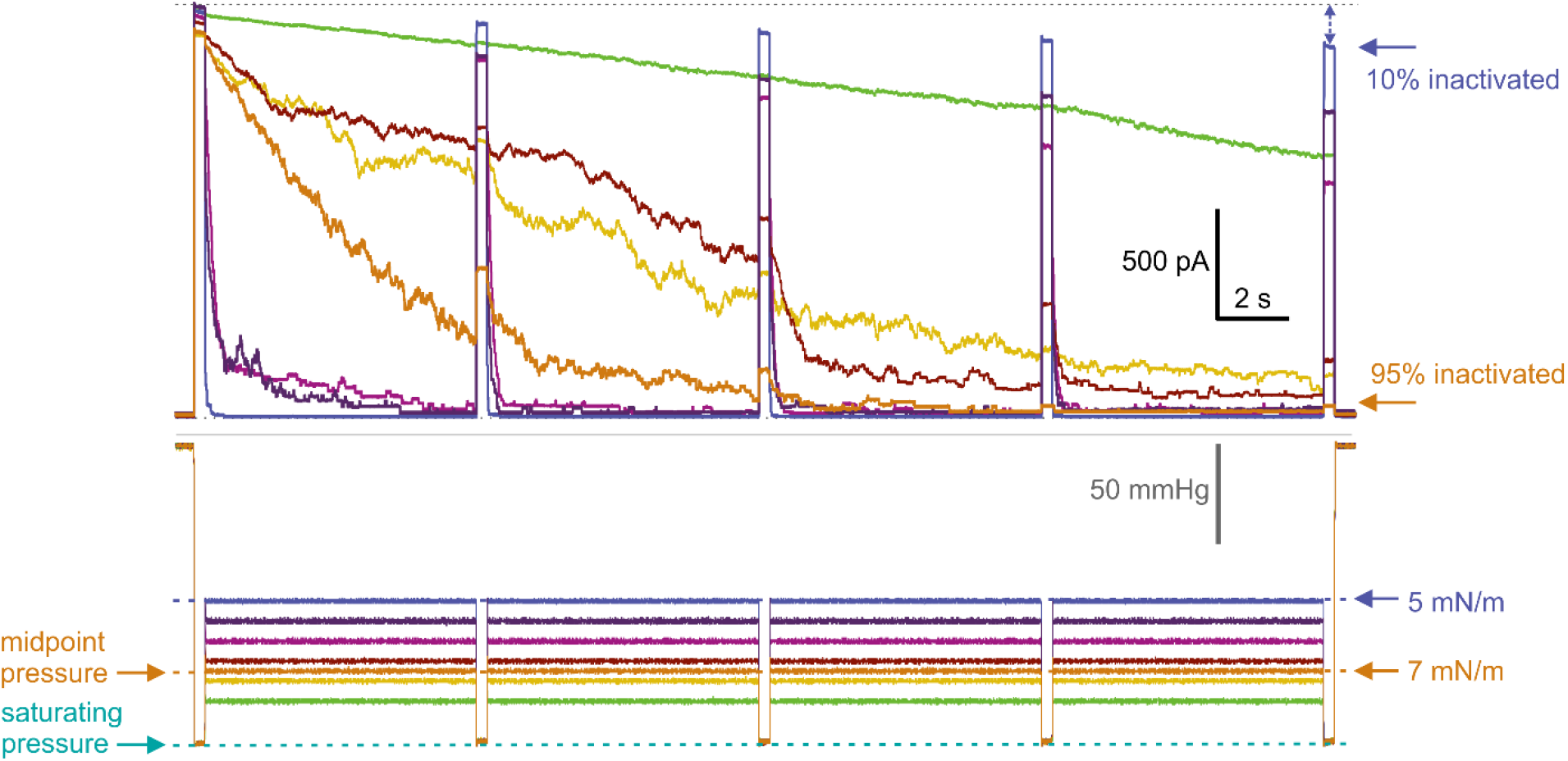
Determination of the tension threshold for inactivation. The ‘comb’ protocols are similar to those in Fig. 1B: evenly spaced saturating test pulses applied on top of a prolonged conditioning step. At the conditioning step (115 mmHg), corresponding to the midpoint pressure (7 mN/m tension), inactivation at the end of the protocol reached a maximum (95%). Increasing the conditioning step amplitude slows adaptation but leads to immediate inactivation (green trace). Decreasing amplitude below the midpoint hastened adaptive closure (magenta, purple, and blue traces) and reduced the extent of inactivation. At 5 mN/m (blue trace), 28% below the midpoint, we still observe 10% inactivation. This illustrates that MscS inactivation has a low-tension threshold.

### MD simulations: the steered transition from the splayed to the flattened state

We started with 6PWP (37), the MscS structure with the complete N-terminus, which was initially equilibrated in a bilayer mimicking an *E. coli*-like mixture (PE/PG, 3:1) for 100 ns. This conformation was converted to the 6VYK structure (27) via a short, steered simulation. Because the 6VYK structure (27) was resolved starting with residue Trp16, we did not apply any restraints to the N-terminal segment from residues 1 through 16, allowing it to adopt its own natural conformation. The backbone of all residues beyond Trp16 was pulled toward the conformation observed in 6VYK during a 5 ns steered simulation. The average force per backbone atom and the time course of RMSD are shown in Fig. 3A. The adjustment was minor, as the transmembrane domains of these structures deviate by only ~2.1 Å RMSD. After this preparatory step, the 6VYK structure was further equilibrated in the bilayer for 100 ns, with the backbone restrained at 1 kcal/mol/Å.

**Fig. 3.**
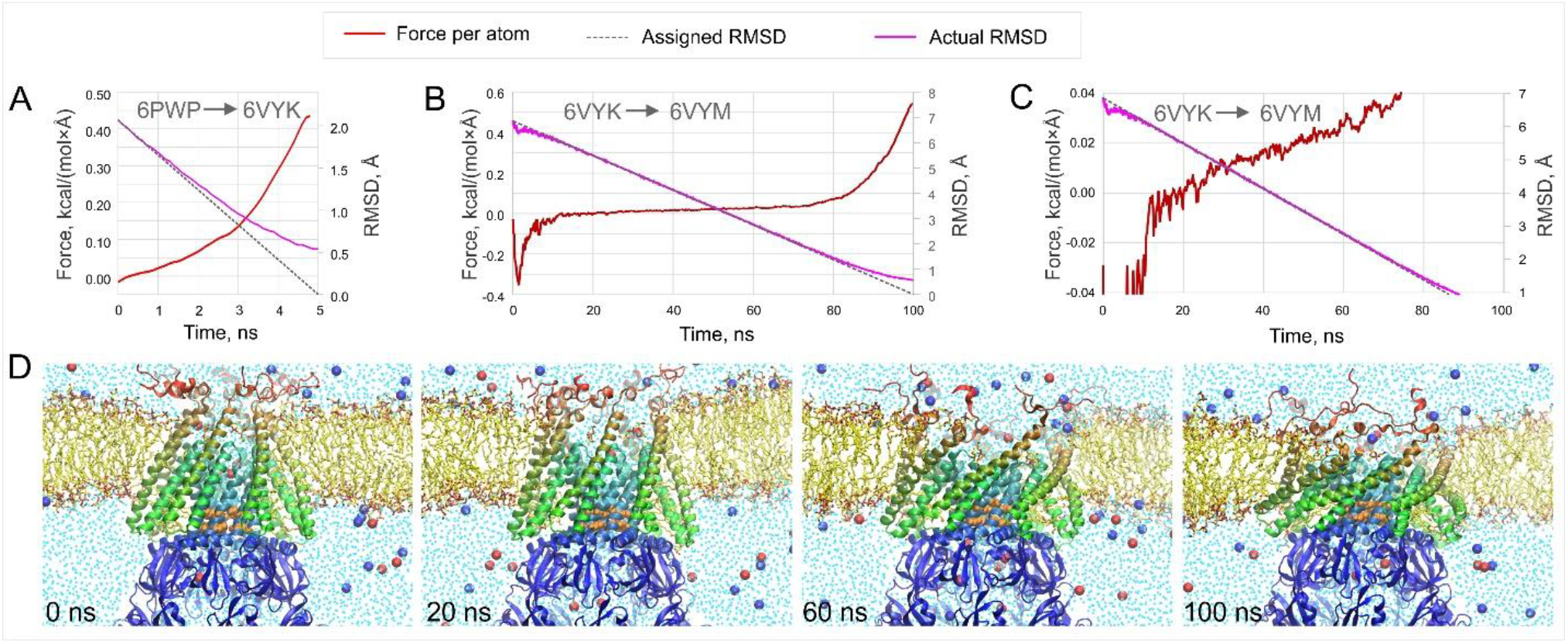
Steered MD simulations of the splayed 6VYK conformation toward the flattened 6VYM state (27). Panels A-C show the time courses of the RMSD change, both the assigned and the actual, as well as the generalized steering force per atom applied only to the backbone. Panel A represents a preparatory step that generated the 6VYK conformation (27) from a more complete 6PWP structure (37). Panel B shows the force profile along the main transition from 6VYK to 6VYM (27) Panel C shows the same view at 10× magnification on the force scale. The monotonous increase in moderate force indicates a barrier-free transition. (D) Snapshots illustrate the sequence of the initial, final, and two intermediate states. The full protein is shown in ribbons, whereas water and lipids are shown within a 10Å vertical slab centered on the protein’s vertical axis.

In the next step, the steering from 6VYK toward the expanded 6VYM conformation occurred within 100 ns, maintaining a linear time course along the RMSD reaction coordinate. This transition mimics the application of extreme tension generated by lipid depletion from nanodiscs with Methyl-β-cyclodextrin (27). The trajectory of the generalized steering force was monotonous and smooth (Fig. 3B). Under the steering protocol, which specifies a linear change in RMSD, the initial steering force per atom was negative, reflecting spontaneous thermal motion of the system, liberated from relatively stiff restraints. Then the average steering force becomes positive and slowly grows in the range between 0 and 0.02 kcal/(mol×Å), signifying an almost frictionless transition, as illustrated in the magnified view (Fig. 3C). After passing 80% of the reaction (from the initial 6.7 to 1.5 Å RMSD), the generalized force began to increase more steeply, reaching 0.55 kcal/(mol×Å) to pull the structure toward the target more forcefully. This final effort is to suppress thermal fluctuations in a system being stiffly restrained to a nearly singular target conformation. Importantly, during the transition, the system encountered no barriers, suggesting smooth conformational connectivity of the two states.

The initial frame, two intermediates, and the final frame illustrate this pathway in Fig. 3D. The structure expanded within the mid-plane of the membrane, increasing the tilt of peripheral helices from ~30° to 67° (27). The structure flattened, resulting in some distortion of the surrounding bilayer. The N-terminal ends that were previously proposed to serve as periplasmic anchors (37), became disordered and lost its tight association with the membrane immediately after the restraints were released.

The characteristic property of the initial 6VYK state (27), as well as similar 6PWP (37), 6RLD (38), 7OO6, 7OO8 (39), 9P0N (26), is the splayed state of the peripheral (TM1-TM2) helical pairs with interhelical gaps filled with lipids in a non-bilayer orientation. These lipids stably occupy the gaps and physically separate the peripheral (tension-receiving) helical pairs from the gate. Interestingly, these lipids partially remain in the structure upon flattening.

The most striking observation was the dramatic flattening of the transmembrane barrel of MscS, without appreciable expansion of the pore. The initial pore diameter between the van der Waals surfaces of the terminal carbons of L105 was 6.1 Å, whereas at the end of the trajectory it was 6.5 Å. The initial pore was largely de-solvated (Figs. 4A and 4D), with occasional water strings present. Based on time courses of water presence (Fig. 4C) and the time-averaged map (Fig. 4E), the flattened 6VYM conformation was slightly more hydrated. Notably, despite a significant electrostatic bias (−200 mV in the cytoplasm), all ions were entirely excluded from the pore constriction. The majority of the ions entering the pore vestibules were Cl^−^, whereas K^+^ rarely appeared in the pore. For the combined 300 ns of simulations, including the equilibrium simulations of both states and the steered transition, no ions crossed the constriction, indicating that both the splayed and the flattened states remained non-conductive. We realize that the steered transition may not precisely reflect the transition experienced by the splayed initial structure upon lipid depletion from the nanodiscs. Yet the smooth pathway did not require extreme forces (Fig. 3) and suggests that the two states are conformationally interconnected, with no substantial barriers in between and no signature of the intervening conductive (open) state proposed in (27).

**Fig. 4.**
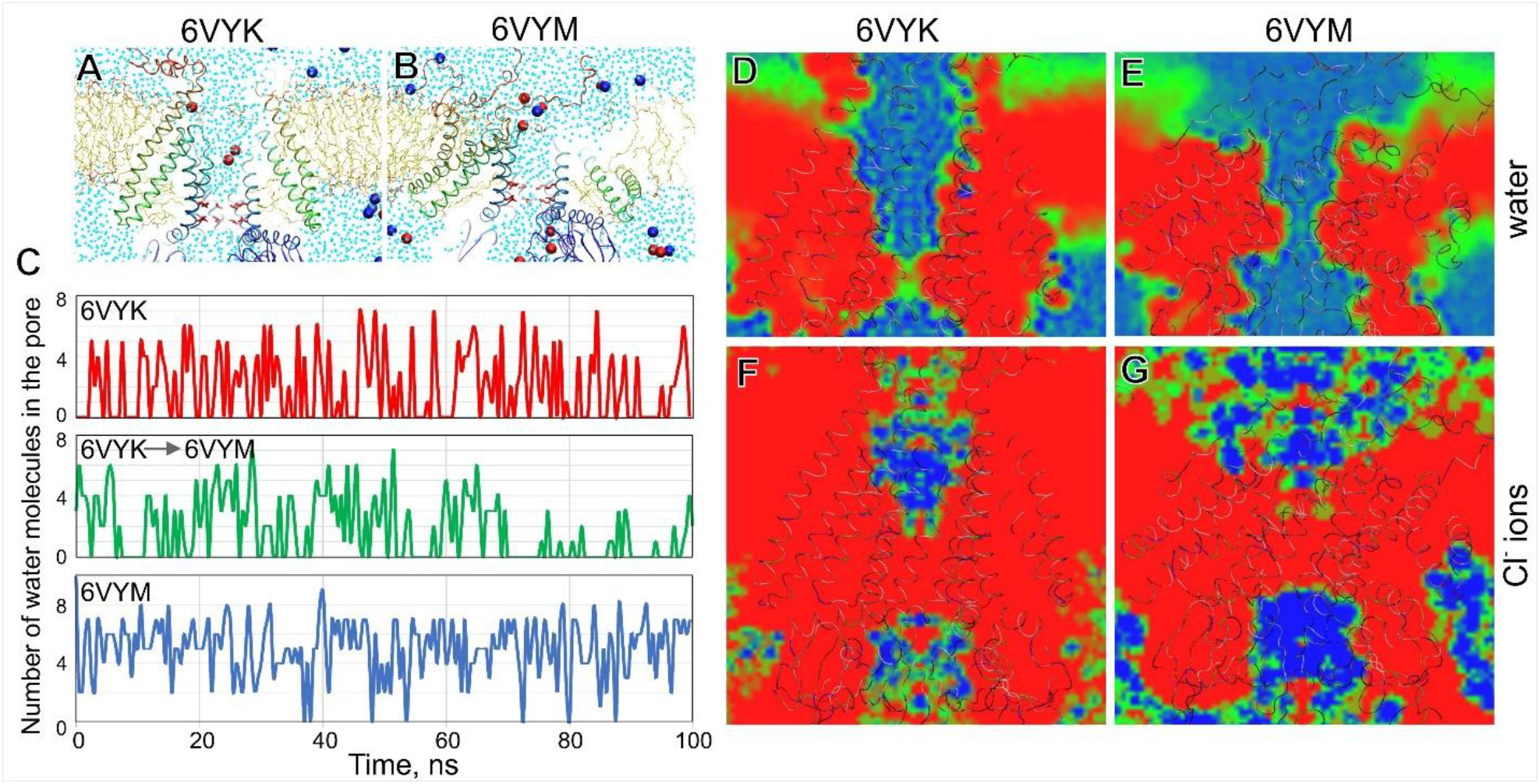
The analysis of water and ion occupancies of MscS pores in the simulated initial 6VYK (27) and final 6VYM (27) conformations, as well as during the transition (6VYK → 6VYM). Snapshots of the initial (A) and final (B) frames of the 100 ns transition. The snapshots are shown as 5A slabs crossing the center of the simulation cell; water is shown as light blue dots; Cl^−^ ions are red, and K^+^ ions are blue. (C) Time courses of water occupancy inside the pore constriction between L105 and L109 during 100-ns equilibrium simulations of 6VYK (red trace), 6VYM (blue trace), and during the transition (green trace). Averaged water density maps averaged over 100 ns equilibrium simulations for the initial (D) and final (E) conformations. Cl^−^ ion density maps for the two conformations (F, G), which show complete exclusion of ions from the pore. All simulations were performed at a near-physiological voltage of −200 mV in the cytoplasm relative to the periplasmic compartment.

## DISCUSSION

The presence of populations of highly conductive osmo-protective channels in the energy-coupling cytoplasmic membranes of essentially all free-living bacteria, as well as in most pathogens and commensals, requires tight regulation. The main question motivating our studies of adaptive MscS behavior is: what if, during active growth and high metabolic activity, the tension in the cytoplasmic membrane fluctuates and occasionally exceeds its low activation threshold? Our data shows that under these conditions, most MscS channels should inactivate. The interval of tensions between MscS activation threshold and midpoint (5-7 mN/m) is precisely the range at which MscS has the highest probability of inactivation (23,25). This must preclude sporadic flickering to the open state at moderate tensions. Once inactivated, the channel cannot be reactivated at any tension unless it is returned to zero for several seconds to recover. A fast, steeply tension-dependent transition to the open state excludes inactivation, whereas tensions lingering at a near-threshold level result in slow, silent inactivation from the (adapted) closed state (23,24,40). The MscS transition to the inactivated state at low tension is sharply accelerated by crowding agents present on the cytoplasmic side (41). The crucial phenomenology of MscS adaptation, inactivation, and recovery, collected using specialized pressure protocols on a single patch, is shown in Fig. 1. The drastic difference in the kinetics of the opening and inactivation transitions is explained by two separate conformational pathways to the open and inactivated states (Fig. 5), both starting from the closed state (24). The fast opening and closing transitions are essentially frictionless (42), whereas the inactivation and recovery processes involve rearrangements of the protein-lipid boundary and thus are much slower (35).

**Fig. 5.**
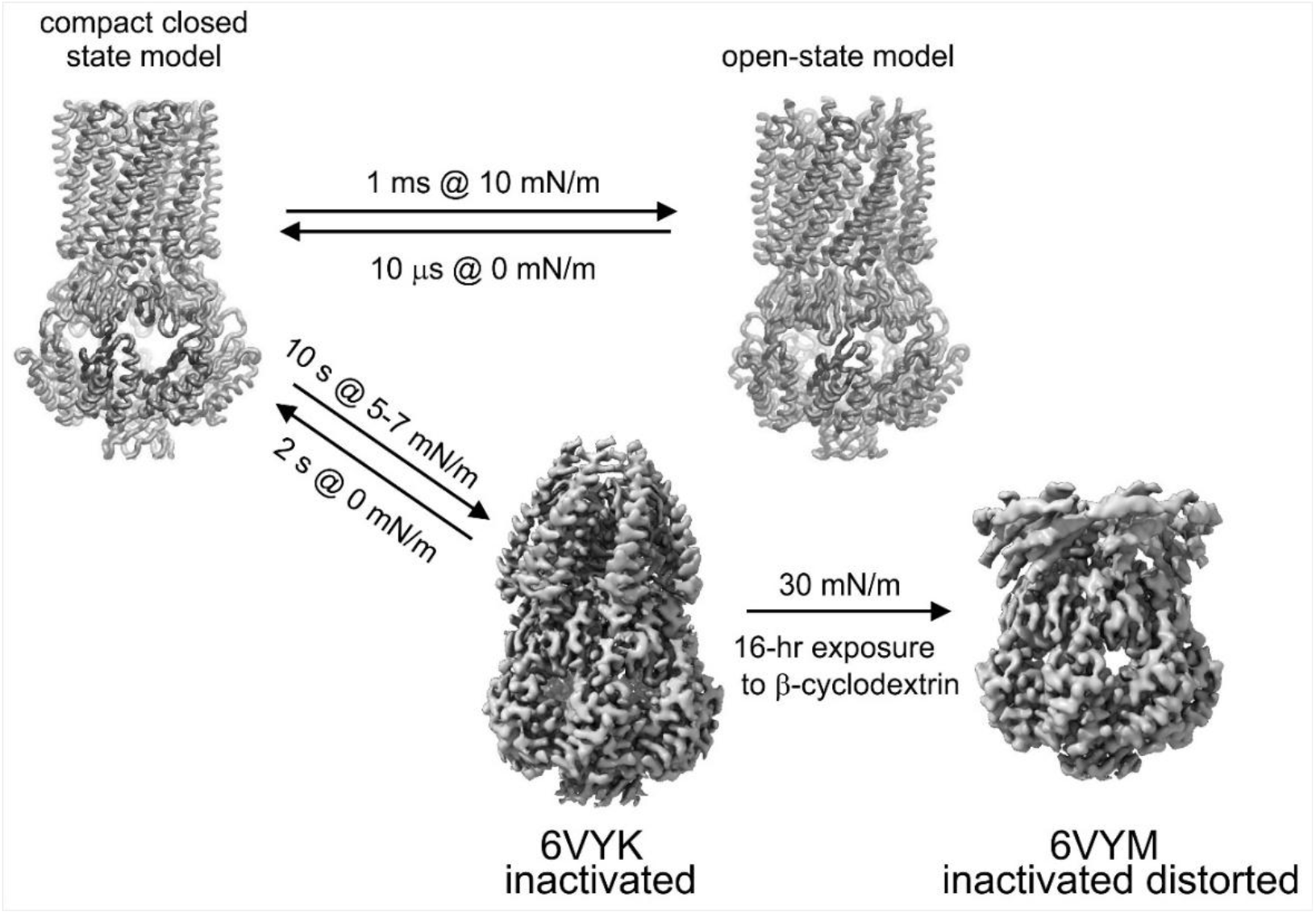
The kinetic scheme of WT MscS with two separate transitions from the closed to open and from the closed to the inactivated state. The functional assignment of the new 6VYK and 6VYM structures (27) expands the inactivation pathway. The opening and inactivation pathways, both starting in the compact closed state, are kinetically separated: the opening and closing transitions can be rapid and involve only minor rearrangements of the protein-lipid boundary, whereas the slow inactivation and recovery transitions apparently involve lipid rearrangement (35). We assign the 6VYK structure (27) as the inactivated state, place the flattened 6VYM structure (27) beyond it, and assign it as a distorted but resilient, non-conductive state.

Despite multiple attempts to rationalize the functional cycle of MscS at the structural level (37,43–45), deep inconsistencies remain. Today, we have two predominant classes of MscS structures: splayed, non-conductive structures similar to the initial crystal structure (15,16), with a characteristic kink that breaks TM3 at G113 (2OAU class), and expanded, partially open states resembling the 2VV5 structure of the A106V mutant (46) (2VV5 class). The splayed conformations are most frequently observed in detergent micelles mixed with regular lipids or in nanodiscs formed with regular lipids (16-18 carbon chains). The expanded 2VV5-like structures with a ~9 Å aqueous pore and partially straightened TM3 helices are observed in either completely delipidated MscS samples (47) or in nanodiscs reconstituted with short-chain lipids (27,37). As soon as a 2VV5-like structure sees regular lipids, it converts into a kinked and splayed non-conductive 2OAU-like state (39).

Zhang and coworkers recently solved the splayed non-conductive conformation of MscS in PC18 nanodiscs (6VYK) and, following that, determined the unique, highly flattened structure of the third type (6VYM), obtained after treatment of these nanodiscs with β-cyclodextrin (27). This treatment generated extreme tension (~30-40 mN/m) inside nanodiscs, which distorted the structure, but did not open the pore. The authors postulated that the initial splayed conformation of WT MscS corresponds to the closed state, which can open under tension. They also assigned the expanded (2VV5-type) conformation commonly observed in delipidated samples (39,47) as open. These postulates have led to expectations that were difficult to prove, turning the paper into a collection of negative yet interesting results. The expectation that the splayed 6VYK conformation (27), when stretched in nanodiscs, would produce a 2VV5-type structure (46) did not hold for either WT MscS or the non-inactivating G113A mutant. The splayed structure flattened under extreme non-physiological tension but did not open. In addition, Zhang and coworkers solved the MscS structure in nanodiscs formed from short-chain (PC10) lipids (6VYL (27)), which belongs to the 2VV5 class of structures (46). This expanded structure had an aqueous pore of about 10 Å and did not seem to satisfy the predicted conductive properties of the open state (48). The exploration of channel behavior in thin bilayers (short-chain-lipid nanodiscs) is interesting as it revealed the MscS barrel deformability, but it is hardly compatible with moderate tensions (5-7 mN/m) that drive all transitions (Fig. 1), at which membrane thinning is not expected. The authors conclude that a hydrophobic (thickness) mismatch alone cannot drive opening (27), and we completely agree.

In their interpretation, the authors followed the typical time sequence of functional states observed in patch-clamp under sub-saturating tensions: closed → open → desensitized. Failing to observe an expanded conductive state in nanodiscs made with regular lipids, they included the 2VV5 structure (46) in the middle of their gating scheme as “open” and assigned the flattened conformation that followed as desensitized. Classifying this conformation as desensitized was a misnomer, because it is (i) contrary to the established definition of the desensitized state as one that can be reactivated at higher tension (23,24), and (ii) disregards the fact that desensitization occurs at moderate sub-saturating tensions (Fig. 1). Note that the flat non-conductive conformation was obtained under extreme 30 mN/m tension and apparently could not be reactivated at any cost.

The linear kinetic scheme terminating in a desensitized state proposed by Zhang and coworkers contradicts the basic phenomenology of MscS gating previously described in (23–25,40) and summarized again in Figs. 1 and 2 of this study. MscS exhibits low-tension transitions to the desensitized and inactivated states and no desensitization or inactivation from the open state at saturating tensions (23). The actual channel behavior is reflected in the gating scheme depicted in Fig. 5, where the compact closed and fully open states were computationally reconstructed (48,49). The forked scheme represents independent tension-driven transitions to the open and inactivated states, both originating from the predicted compact closed state. In this scheme, the splayed 6VYK conformation, with lipid-facing TM1-TM2 helices separated from the gate, is designated as the inactivated state. Although not shown in the published density map (Electron Microscopy Data Bank EMD-21462), the illustrations in the main text (27) show that the gaps between the peripheral helical pairs (TM1-TM2) in 6VYK are filled with non-bilayer lipids, apparently stabilizing the uncoupled state of the gate. The details of lipid packing in MscS crevices, which physically separate TM2 from TM3, were recently reported in a structure by Moller et al., (PDB 9P0N (26)). The kinetic scheme (Fig. 5) predicts that at zero tension, part of the channel population recovers to the closed state, designated as a ‘compact closed model’ in which TM1-TM2 pairs are aligned with TM3 and reconnected to the gate. Abruptly applied super-threshold tension to the closed conformation results in opening, accompanied by in-plane protein expansion of 18-20 nm^2^ and the formation of a ~16 Å aqueous pore (23,35,48). A prolonged action of low tension, instead, leads to silent inactivation, which is associated with in-plane expansion of 8-9 nm^2^ due to lipid penetration into interhelical crevices and helical splay (24).

Our simulations show that the 6VYK structure is completely non-conductive (Fig. 4), and it did not open under extreme tension, forming the flat 6VYM conformation (27), also non-conductive. The steered transition between the two states indicated conformational connectivity between them without transitioning through a conductive state. This strongly suggests that 6VYK is an unperturbed inactivated state, whereas 6VYM is a substantially distorted inactivated state that resists opening under extreme tension (27), directly supporting our results in Fig. 1D.

We re-assign the class of splayed conformations not as a closed but as an inactivated state. We find the structural information in Zhang’s paper highly instructive and perfectly matches the functional behavior of MscS presented above. Inactivated MscS cannot be opened at any tension attainable in patch-clamp, nor can it be opened in nanodiscs under any tension. This is highly consistent with multiple MD simulations showing that the splayed conformations could not be substantially opened by applying tension to lipids surrounding MscS in simulated cells (44,45,50,51). Our analysis places the flattened 6VYM structure (27) as an extreme of the inactivation path on the right side of the functional diagram (Fig. 5).

Having a large population of easily activated (low-threshold) mechanosensitive channels in the cell’s cytoplasmic membrane that maintains vital gradients would be counterproductive. However, these channels are necessary to protect the cells osmotically. Therefore, the MscS’s ability to inactivate is essential for bacterial cells’ metabolic efficiency, competitiveness, and survival. Strains with non-inactivating MscS mutants show toxic ‘gain-of-function’ phenotypes and cannot properly terminate the large permeability response in the event of strong osmotic shock (2). Inactivation and tight regulation allow large populations of low-threshold mechanosensitive channels to be present safely in the energy-coupling membrane. When inactivated, MscS resists opening, helping preserve the membrane’s integrity and barrier function.

## ACKNOWLEGEMENTS

We thank Samantha Miller for providing the MJF465 strain. This work was supported by NIH R01-AI135015 to SS. This material is based upon work supported by the National Science Foundation Graduate Research Fellowship Program to EM under Grant No. DGE 1840340. Any opinions, findings, and conclusions or recommendations expressed in this material are those of the author(s) and do not necessarily reflect the views of the National Science Foundation.

## AUTHOR CONTRIBUTIONS

AA: molecular dynamics simulations, data interpretation, figure making, writing

EM: patch-clamp electrophysiology, data interpretation, figure making, writing, funding

SS: data interpretation, writing, supervision and training, funding

## DATA AVAILABILITY

The data that support this study are available from the corresponding authors upon request.

## CODE AVAILABILITY

The scripts used in this study are available from the corresponding authors upon request.

